# The essential *Porphyromonas gingivalis* type IX secretion system component PorZ delivers anionic-lipopolysaccharide to the PorU sortase for transpeptidase processing of cargos

**DOI:** 10.1101/2020.07.16.206169

**Authors:** Mariusz Madej, Zuzanna Nowakowska, Miroslaw Ksiazek, Anna M. Lasica, Danuta Mizgalska, Magdalena Nowak, Anna Jacula, Carsten Scavenius, Jan J. Enghild, Joseph Aduse-Opoku, Michael A. Curtis, F. Xavier Gomis-Rüth, Jan Potempa

## Abstract

Cargo proteins of the type IX secretion system (T9SS) in human pathogens from phylum Bacteroidetes invariably possess a conserved C-terminal domain (CTD) that functions as a signal for outer membrane (OM) translocation. In *Porphyromonas gingivalis*, the CTD of selected cargos is cleaved off after translocation, and anionic lipopolysaccharide (A-LPS) is attached. This transpeptidase reaction anchors secreted proteins to the OM. PorZ, a cell surface-associated protein, is an essential component of the T9SS whose function was previously unknown. We recently solved the crystal structure of PorZ, and found that it consists of two β-propeller moieties followed by a CTD. In this study, we performed structure-based modelling suggesting that PorZ is a carbohydrate-binding protein. We found that recombinant PorZ specifically binds A-LPS. Binding was blocked by monoclonal antibodies that specifically react with a phosphorylated branched mannan in the anionic polysaccharide (A-PS) component of the A-LPS, but not with the core oligosaccharide or the lipid A endotoxin. Examination of A-LPS derived from a cohort of mutants producing various truncations of A-PS confirmed that the phosphorylated branched mannan is indeed the PorZ ligand. Moreover, purified recombinant PorZ interacted with the PorU sortase in an A-LPS–dependent manner. This interaction on the cell surface is crucial for the function of the attachment complex composed of PorU, PorZ, and the integral OM β-barrel proteins PorV and PorQ, which is involved in post-translational modification and retention of T9SS cargos on the bacterial surface.

**Author summary:** Bacteria have evolved multiple systems to transport effector proteins to their surface or into the surrounding milieu. These proteins have a wide range of functions, including attachment, motility, nutrient acquisition, and toxicity in the host. *Porphyromonas gingivalis*, the human pathogen responsible for severe gum diseases (periodontitis), uses a recently characterized type IX secretion system (T9SS) to translocate and anchor secreted virulence effectors to the cell surface. Anchorage is facilitated by sortase, an enzyme that covalently attaches T9SS cargo proteins to a unique anionic lipopolysaccharide (A-LPS) moiety of *P. gingivalis*. Here, we show that the T9SS component PorZ interacts with sortase and specifically binds A-LPS. Binding is mediated by a phosphorylated branched mannan repeat in A-LPS polysaccharide. A-LPS– bound PorZ interacts with sortase with significantly greater affinity, facilitating modification of cargo proteins by the cell-surface attachment complex of the T9SS.

## Introduction

Periodontal disease is one of the most prevalent infection-driven chronic inflammatory diseases in humans [1]. The disease is the result of a host immune response against dysbiotic microorganisms accumulated in the gingival crevice that results in destruction of tooth-supporting tissues. If left untreated, up to 15% of affected adults develop severe periodontitis, which may result in tooth loss [2]. In addition, periodontal infection is associated with systemic disorders such as Alzheimer’s disease, osteoporosis, diabetes, rheumatoid arthritis, and respiratory and cardiovascular diseases [3–6].

The Gram-negative anaerobic bacterium *Porphyromonas gingivalis* is a major periodontal pathogen that resides in the human gingival crevice. As an asaccharolytic organism, it acquires nutrients in the form of peptides from proteins localized in the gingival crevicular fluid [7,8]. In contrast to the majority of Gram-negative bacteria, *P. gingivalis* produces two different types of lipopolysaccharide (LPS) molecules [9–11]. Both contain the conserved lipid A endotoxin and a core oligosaccharide, but they differ in the highly variable polysaccharide (PS), called the O-antigen, attached to the core. The PS of the more common O-LPS (O-PS) consists of repeating units of a single tetrasaccharide, whereas A-LPS consists of a phosphorylated branched mannan (anionic polysaccharide, A-PS) [9–11] (Fig 1).

**Fig 1.**
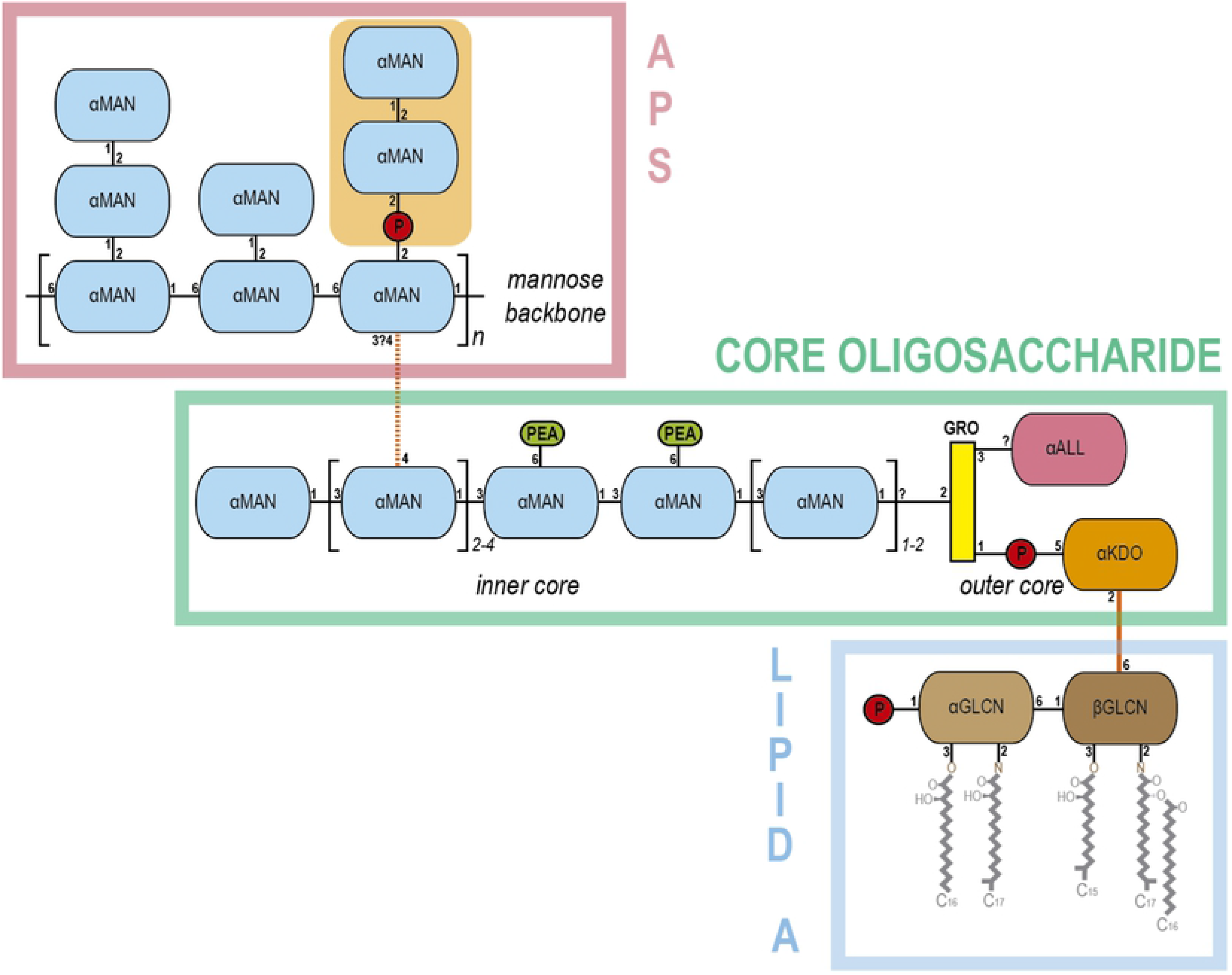
Anionic lipopolysaccharide (A-LPS) of *Porphyromonas gingivalis*. Scheme depicting the modular structure of the anionic lipopolysaccharide (A-LPS) of *P. gingivalis.* The A-LPS consists of the A-type anionic distal polysaccharide (APS), the core oligosaccharide, and the endotoxin or lipid A. The APS consists of an α-D-mannose (αMAN) backbone, which has side chains of one or two αMANs. One of these side chains contains a branched phosphomannan element (phosphate groups are shown as red circles, orange background). The core oligosaccharide spans a linear chain of αMANs that constitute the inner core. One of the monomers is linked to the mannose backbone of APS (orange dashed line). The two central αMANs of the inner core are linked to phosphoethanolamine (PEA). The inner core chain of αMANs is linked to the outer core through a glycerol molecule (GRO). The latter is linked to α-D-allosamine (αALL) and to a 3-deoxy-D-α-manno-oct-2-ulopyranosonic acid (αKDO) via a phosphate. Monomer αKDO is linked (orange line) to a β-D-glucosamine (βGLCN) from lipid A. The latter is linked to a α-D-glucosamine (αGLCN) and gives rise to the disaccharide bone structure of lipid A, which carries a phosphate and the lipids.

To thrive in the inflammatory environment of its colonization site [12], *P. gingivalis* secretes an array of proteinaceous virulence factors. These proteins, including the major extracellular proteolytic agents RgpA, RgpB, and Kgp (collectively called gingipains) [13,14], possess an N-terminal signal peptide that directs their translocation through the inner membrane (IM) into the periplasmic space via the *Sec* system. In addition, they have a conserved C-terminal domain (CTD) with an Ig-like fold involved in targeted transport across the outer membrane (OM) via the type IX secretion system (T9SS) [15–19].

Despite participating only in OM translocation, the T9SS spans both the IM and the OM [20–22]. Its protein components reside on the IM (PorL and PorM) [23,24] or in the periplasm (PorN, PorK, PorW, and PorE) [25–27]; constitute the OM translocon itself (Sov, PorV, PIP, and Plug) [28]; or are anchored to the cell surface (PorZ and PorU) via interactions with integral OM β-barrel proteins (PorV and PorQ, respectively) [29]. After translocation through the OM, the surface-located PorV-anchored sortase PorU cleaves the CTD and attaches an anionic lipopolysaccharide (A-LPS) molecule, thereby fastening secreted proteins to the OM [30]. Expression of the major T9SS components is regulated by PorX and PorY, which form a unique two-component system [31].

Inactivation of any essential component of the T9SS, including PorZ [32], leads to lack of pigmentation in the mutant strain and accumulation of unprocessed proteins, e.g., inactive progingipains, in the periplasm. In this study, we found that in addition to progingipains, ΔPorZ also accumulates A-LPS in the periplasmic space, as previously described for *porV*^−^(ΔPorV), *porT*^−^ (ΔPorT), and *porU*^−^ (ΔPorU) deletion mutants [33]. Furthermore, we showed that PorZ interacts with A-LPS through the phosphorylated branched mannan (Manα1-2Manα1-phosphate) of the A-PS. Finally, we found that A-LPS promotes the interaction between PorZ and the PorU sortase, supporting the idea that together with PorV and PorQ, PorZ and PorU form a large attachment complex engaged in modifying and anchoring T9SS cargos to the *P. gingivalis* surface [29].

## Methods

### Bacterial strains and general growth conditions

*P. gingivalis* strains (wild-types W83 and HG66 and mutants; listed in **S1 Table**) were grown in enriched tryptic soy broth (eTSB) (30 g trypticase soy broth and 5 g yeast extract per litre at pH 7.5; further supplemented with 5 mg hemin, 0.5 g L-cysteine, and 2 mg menadione) or on eTSB blood agar (eTSB medium containing 1.5% [w/v] agar; further supplemented with 4% defibrinated sheep blood) at 37°C in an anaerobic chamber (Don Whitley Scientific, UK) with an atmosphere of 90% nitrogen, 5% carbon dioxide, and 5% hydrogen. *Escherichia coli* strains (listed in **S2 Table**), used for all plasmid manipulations, were grown in Luria–Bertani (LB) medium and on 1.5% LB agar plates. For antibiotic selection in *E. coli*, ampicillin was used at 100 μg/ml. *P. gingivalis* mutants were grown in the presence of erythromycin at 5 μg/ml and/or tetracycline at 1 μg/ml.

### Generation of ΔPorZ, ΔPorT, and ΔPorV deletion mutants of *P. gingivalis*

The *P. gingivalis* deletion mutants ΔPorZ and ΔPorT were generated by homologous recombination, as previously described [32, 34]. ΔPorV was created using the same strategy that was previously reported for ΔPorZ and ΔPorT. In brief, two 1-kb segments of DNA flanking the *porV* gene in *P. gingivalis* W83 (TIGR accession number PG0027) were amplified by PCR using Accuprime Pfx DNA polymerase (Invitrogen) and primer pairs PG23FrBXbaIF+PG23FrBHind3R and PG23FrASacIF+PG23FrASmaR (sequence listed in **S3 Table**) and ligated into the multiple cloning site of the pUC19 plasmid at XbaI+HindIII and SacI+SmaI sites, respectively. An erythromycin-resistance cassette, *ermF*-*ermAM*, was amplified by PCR from pVA2198 [35] using the primer pair ermFAMSmaIF/ermFAMXbaIR and ligated between the flanking DNA segments at SmaI and XbaI sites to create the final plasmid, p23AeB-B. One μg of p23AeB-B was electroporated into electrocompetent *P. gingivalis* cells to allow for homologous recombination of the construct into the genome to result in deletion of *porV* and acquisition of the *ermF*-*ermAM* cassette. Erythromycin-resistant *P. gingivalis* colonies were subsequently confirmed for deletion of *porV* via PCR amplification and sequencing of the pertinent region of the genome. Deletion mutants of PG0129 (α-1,3-mannosyltransferase) and PG1142 (Wzy, O-antigen polymerase) were constructed and characterised by Rangarajan and colleagues [36]. Deletion of PG1142 leads to a core-plus-one repeating unit structure for both LPS types, whereas inactivation of PG0129 leads to the absence of A-PS and O-PS moieties.

### Site-directed mutagenesis of the porZ of *P. gingivalis*

All point mutations (K42A, R132A, and R338A; numbering according to UniProt database entry Q9S3Q8) were created on master plasmids (p1604CeB-H and pAL33; described below) using the Phusion Site-Directed Mutagenesis Kit (Thermo Scientific) followed by electroporation into wild-type *P. gingivalis* for homologous recombination [37]. Newly generated plasmids and *P. gingivalis* mutants were confirmed by sequencing (CGeMM DNA Facility Core, University of Louisville, KY, USA). For primer names and sequences, see **S3 Table**.

A single mutation, R338A, of PorZ was created on the p1604CeB-H master plasmid [32]. The plasmid contains the erythromycin-resistance cassette (*ermF*-*ermAM*) flanked at 5’ with the whole open reading frame of *porZ* (PG_RS07070/PG1604) and at 3’ with a DNA fragment present downstream of *porZ*. The final mutagenesis plasmid was named pAL31 and used to create a *P. gingivalis* PorZ^R338A^ strain (AL008).

Mutations of K42A and R132A in PorZ were created on the pAL33 master plasmid. This plasmid is a pUC19 derivative, which contains a fragment of the *porZ* gene (1–1248 bp/416 aa) cloned together with its upstream DNA fragment (861 bp). The primer pair used for cloning (**S3 Table**) overlaps with the naturally occurring restriction sites NdeI (upstream DNA) and EcoRI (within the *porZ* gene). After introducing point mutations, the transient constructs pAL34 (R132A) and pAL36 (K42A) were obtained. In the next step, mutated *P. gingivalis* DNA was excised using NdeI and EcoRI and ligated with p1604CeB-H digested with the same restriction enzymes to obtain final plasmids pAL39 (R132A) and pAL40 (K42A). They have a structure similar to p1604CeB-H but contain additional DNA upstream of *porZ*, which was sufficient enough for homologous recombination to introduce mutations at the beginning of the *porZ* gene (see **S2 Fig** for a schematic view). Final mutagenesis plasmids were used to create *P. gingivalis* strains expressing PorZ^K42A^ and PorZ^R132A^, referred to as AL015 and AL016, respectively. In the same manner, double PorZ^K42A/R338A^ (AL017) and PorZ^K42A/R132A/R338A^ (AL018) mutants were created. For a full plasmid description, see **S2 Table**.

### Gingipain activity assay

RgpA/B and Kgp protease activities of *P. gingivalis* were determined as described previously [38] using the chromogenic *p*-nitroanilide (*p*NA) substrates benzoyl-Arg-*p*NA and acetyl-Lys-*p*NA, respectively. The rate of substrate hydrolysis was recorded as the increase in optical density (OD) at 405 nm due to the release of *p*NA at 405 nm (SpectroMax M5 microplate reader, Molecular Devices). The experiment was repeated twice (in triplicates), and the activity of each enzyme is given as a percentage of wild-type activity.

### Subcellular fractionation of *P. gingivalis* strains

Stationary phase cultures of wild-type *P. gingivalis* and mutants were adjusted to OD_600_ = 1.0, and cells were collected by centrifugation (8,000 × g, 15 min). The cell pellet was then washed and resuspended in PBS. This fraction is referred to as “washed cells” (WC). The collected cell free culture medium was ultracentrifuged (100,000 × g, 1 h) to remove vesicles, and the supernatant was concentrated 10-fold by ultrafiltration using 3-kDa cut-off centricones (EMD, Millipore, Billerica, MA); this fraction was designated as the “clarified medium” (CF). The WC fraction, obtained as described above, was suspended in buffer containing 0.25 M sucrose and 30 mM Tris·HCl at pH 7.6. After a 10 min incubation period, cells were pelleted (12,500 × g, 15 min) and rapidly resuspended in 2.5 ml of cold distilled water to disrupt the OM. After an additional 10 min incubation period, spheroplasts were pelleted by centrifugation (12,500 × g, 15 min). The supernatant was collected and designated as the “periplasmic” (PP) fraction. All fractions were supplemented with peptidase inhibitors (5 mM tosyl-L-lysyl-chloromethyl ketone [TLCK], 1 mM 2,2’-dithiodipyridine [DTDP], 1× EDTA-free protein inhibitor cocktail; all from Roche) before storage at −20°C.

### Western blot analysis

Samples of the washed cells, PP fraction, and medium fraction were resolved by SDS-PAGE and transferred to a nitrocellulose membrane. To visualise the total protein transferred, Ponceau S staining was used. After washing out Ponceau S with H_2_O, membranes were blocked with 5% non-fat skim milk in TBST (0.1% [v/v] Tween-20 in 20 mM Tris (pH 7.5), 500 mM sodium chloride) overnight at 20°C, incubated with primary antibody 1B5 (1 μg/ml) for 2 h at room temperature, followed by incubation with goat anti-mouse horseradish peroxidase-conjugated IgG (1/20,000 dilution) in blocking solution for 1 h at RT. Development was carried out using the ECL Western Blotting substrate kit according to the manufacturer’s instructions (Pierce, UK).

### Digestion of *P. gingivalis* proteins in whole cell extract and the periplasmic fraction by proteinase K

The aliquots of the PP fraction and whole cell extract were heated at 60°C for 1 h. Then, the samples were cooled on ice, and proteinase K (Promega) was added to a final concentration of 100 μg/ml. The samples were incubated at 37°C overnight and then subjected to Western blot with mAb 1B5.

### Isolation of anionic lipopolysaccharide (A-LPS) and total lipopolysaccharide (LPS)

Isolation of A-LPS from the ΔPorV mutant and W83 strain was performed according to a previously described procedure [10]. For affinity chromatography, ion-exchange chromatography, and size-exclusion chromatography, HiTrap ConA 4B, HiPrep DEAE FF 16/10, and Superdex200 Increase 10/300 GL columns, respectively, were used. All columns were purchased from GE Healthcare Life Sciences. Isolation of total LPS from HG66, ΔPG0129, and ΔPG1142 strains was carried out using an LPS extraction kit (Intron Biotechnology). *E. coli* LPS was purchased from Sigma. The concentration of LPS was determined using the Limulus Amebocyte Lysate (LAL) kit (Lonza).

### Flow cytometry analysis

To reduce the low background of mAb 1B5 on HG66 cells, the antibodies at a concentration of 200 μg/ml were first incubated with PBS-washed HG66 cells (OD_600_=0.2) for 1 h at room temperature. The cells were then centrifuged (6000 × g, 15 min), and the supernatant, devoid of non-specific antibodies, was filtered through a 0.2 μm syringe filter. The concentration of antibodies was adjusted to 40 μg/ml and was used for flow cytometry analysis. *P. gingivalis* strains were grown in eTSB until they reached the late exponential stationary growth phase (OD_600_=1.2–1.5). Bacterial cells were harvested by centrifugation, washed twice with PBS, and adjusted to OD_600_=1.0 with staining buffer (PBS supplemented with 2.5% bovine serum albumin). Then, 100 μl of the cell suspension was transferred to a 96-well conical plate and incubated for 30 min with staining buffer. Cells were collected by centrifugation (500 × g, 5 min), and the pellet was resuspended in the staining buffer containing mAb 1B5 at a total protein concentration of 40 μg/ml and incubated for 30 min. Thereafter, cells were centrifuged (500 × g, 5 min) and the newly obtained pellet was resuspended in the staining buffer containing goat anti-mouse antibody conjugated with fluorescein isothiocyanate (Abcam) at a 1:200 dilution and incubated for 30 min. Cells were washed twice with PBS after each incubation with antibodies. The whole staining procedure was performed on ice. After staining, one-color flow cytometry analyses were performed using a FACSCalibur apparatus (BD Biosciences) operating with CellQuest software (BD Biosciences). Graphs were prepared using the FLOWJO v.10.6.2 program (Ashland, USA).

### Expression and purification of PorZ and PorU

PorZ was expressed and purified as previously described [32]. PorU was expressed and purified as reported elsewhere (Mizgalska *et al.*, manuscript in preparation). Protein concentrations were determined by measurement of absorbance at 280 nm using a NanoDrop device (Thermo Fisher Scientific) using a theoretical Abs 0.1% value (1.01) calculated by ProtParam (http://web.expasy.org/).

### Analysis of the interaction between A-LPS and PorZ by microscale thermophoresis

Periplasmic fractions of W83, HG66, ΔPorZ, ΔPorV, and ΔPorT strains were prepared as described earlier. The fractions were assayed for the LPS concentration using the Limulus Amebocyte Lysate (LAL) kit (Lonza) and adjusted to the same LPS content. The PP fractions were treated overnight with proteinase K and then ultrafiltrated with filter devices with a 3 kDa cut-off (Merck Millipore). Proteinase K was then thermally inactivated. A Monolith NT.115 instrument (NanoTemper Technologies GmbH, Munich, Germany) was used to analyse the binding interactions between the recombinant PorZ and the LPS purified from the periplasmic space. PorZ was fluorescently labelled on lysine residues with NT-647 NHS dye. The PP fraction with LPS concentrations ranging from 3 × 10^3^ to 1 × 10^8^ EU/ml was incubated for 5 min at 20°C with 50 nM PorZ. Experiments were performed in PBS at pH 7.4. The samples were loaded into the Monolith NT.115 Standard Treated glass capillaries and initial fluorescence measurements followed by thermophoresis measurements were carried out using 40% LED power and Medium MST power. *K_d_* values were calculated using the MO.Affinity Analysis software. Experiments were performed in triplicate. Binding experiments with purified A-LPS from the ΔPorV mutant and the W83 wild-type strain, commercially available LPS from *E. coli*, and total LPS isolated from *P. gingivalis* HG66, ΔPG0129, and ΔPG1142 were performed analogically. The concentration of LPS used was in the range of 3 × 10^1^ to 1 × 10^5^ EU/ml.

### Analysis of the interaction between PorZ and PorU by microscale thermophoresis

A Monolith NT.115 was used to measure the binding interactions between PorU and PorZ or PorZ saturated with A-LPS from the W83 strain. PorU at a concentration of 3 nM to 100 μM was incubated for 5 min with 20 nM PorZ in PBS. Alternatively, A-LPS from W83 at a concentration of 1×10^5^ EU/ml was added to PorZ, which was then incubated with PorU. The samples were loaded into the Monolith NT.115 Premium Treated glass capillaries, and initial fluorescence measurements were carried out using 60% LED power. A concentration-dependent increase in fluorescence of NT-647-labelled PorZ in the serial dilution of PorU was observed. Therefore, a “standard deviation (SD) test” was performed to discriminate between binding-specific fluorescence quenching and loss of fluorescence due to aggregation of PorZ upon addition of PorU. To this end, samples corresponding to low and high PorU concentrations (3, 6, and 12 nM and 25, 50, and 100 μM, respectively) were first centrifuged for 10 min at 16,000 × *g* to remove protein aggregates and then mixed with a 2× solution containing 4% SDS and 40 nM DTT and heated to 95°C for 5 min to denature PorZ. Subsequently, the fluorescence of these samples was measured using a Monolith NT.115 instrument. If the increase in fluorescence in the serial dilution is binding induced, the fluorescence intensities after denaturation should be identical, independent of the titrant concentration, which was the case for PorU-PorZ interactions regardless of the presence of A-LPS. *K*_*d*_ values were calculated using the MO.Affinity Analysis software. Experiments were performed in triplicate.

### Co-purification assay

The purified PorZ and hexahistidine-tagged PorU were mixed in equimolar concentration and incubated on ice for 10 min. The mixture was then applied to Co^2+^- nitrilotriacetic acid (Co^2+^- NTA) magnetic beads (Novex) in 100 mM sodium phosphate (pH 8.0), 600 mM sodium chloride, and 0.02% Tween-20 and incubated at 25°C for 10 min. The beads were washed three times with 100 mM sodium phosphate (pH 8.0), 600 mM sodium chloride, and 0.02% Tween-20 supplemented with 10 mM imidazole and eluted with 50 mM sodium phosphate (pH 8.0), 300 mM sodium chloride, 0.01% Tween-20, and 300 mM imidazole. Fractions were subjected to SDS-PAGE.

## Results

### A-LPS accumulates in the periplasm of the ΔPorZ mutant

Cargo proteins of the T9SS are anchored in the outer layer of the OM via covalently attached A-LPS. This type of LPS accumulates in the periplasm in the ΔPorV mutant [33].To determine whether this is a general feature of secretion mutants that ablate translocation, we used the monoclonal antibody 1B5 (mAb 1B5), which exclusively recognizes a phosphorylated branched mannan (Manα1-2Manα1-phosphate) of A-LPS [10], to detect the A-LPS in washed cells, isolated periplasm, and clarified growth medium from ΔPorZ and ΔPorT mutants, as well as their parental wild-type W83 strain. ΔPorV fractions were used as positive controls, and strain HG66, which lacks A-LPS but produces intact O-LPS [39], was used as the negative control Western blot analysis revealed that all strains except HG66 had A-LPS in whole-cell extract (Fig 2, WC). By stark contrast, the A-LPS was detected at high levels only in periplasm derived from secretion mutants, but not in periplasm from the wild-type W83 strain (Fig 2, PP). Conversely, the A-LPS was detected exclusively in medium from W83 (Fig 2, CF). Notably, A-LPS moieties in the periplasm of the secretion mutants had lower molecular masses than those in whole-cell extract of the wild type (Fig 2, WC and PP). This observation suggests that protein-depleted A-LPS of 35–40 kDa accumulated exclusively in the periplasm of the mutant strains. The lack of a protein component in this fraction was confirmed by treatment with proteinase K, which did not affect the SDS-PAGE mobility of the A-LPS (S1A Fig). Conversely, when the whole-cell fraction from W83 was treated overnight with proteinase K, the high-molecular weight bands disappeared, confirming that at least some A-LPS in W83 is conjugated to proteins (S1B Fig). Finally, as expected, we observed no mAb 1B5-reactive material in any fraction derived from strain HG66 (Fig 2, WC, PP, CF)). Together, these data support the idea that secretion mutants retain fairly homogenous, protein-unconjugated A-LPS in the periplasm, whereas in the wild-type strain W83, proteins are attached to the OM via A-LPS molecules.

**Fig 2.**
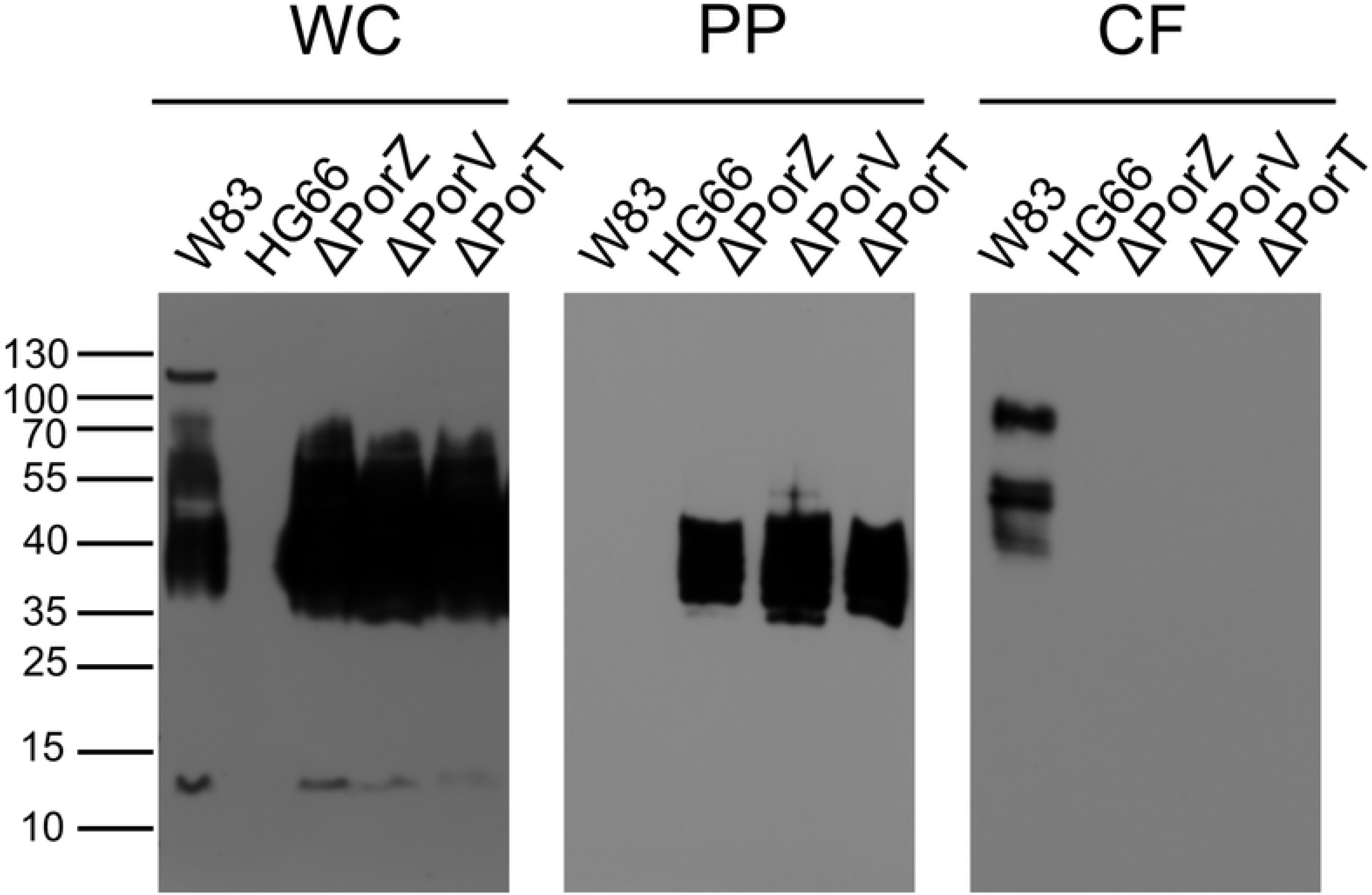
Western blot analysis to detect A-LPS in subcellular fractions of *Porphyromonas gingivalis* cultures. *P. gingivalis* cultures of W83, HG66, ΔPorZ, ΔPorV, and ΔPorT strains were fractionated as described in Methods. Washed cells (WC), periplasm (PP), and clarified medium (CF) from each strain were resolved by SDS-PAGE, transferred onto nitrocellulose membranes, and incubated with mAb 1B5. The medium fraction was concentrated 10-fold before loading on SDS-PAGE.

### The type IX secretion system (T9SS) does not transport the A-LPS to the outer membrane

Periplasmic accumulation of the A-LPS may affect the amount of A-LPS in the cell envelope. Therefore, to quantitatively compare the level of A-LPS in the OM between the secretion mutants and the wild-type strain, we analysed live bacterial cells by flow cytometry with mAb 1B5 (Fig 3). The specificity of the analysis was verified by the absence of staining in strain HG66 (Fig 3F). By contrast, cells of the W83, ΔPorV, and ΔPorT strains reacted strongly with the antibody, with no clear difference in the abundance of the A-LPS among the three (Fig 3B– E). This result indicates that inactivation of genes essential for T9SS function has no significant impact on the level of A-LPS in the OM, implying that the A-LPS is exported in a T9SS independent manner, possibly by some variant of the MsbA/Lpt system [40].

**Fig 3.**
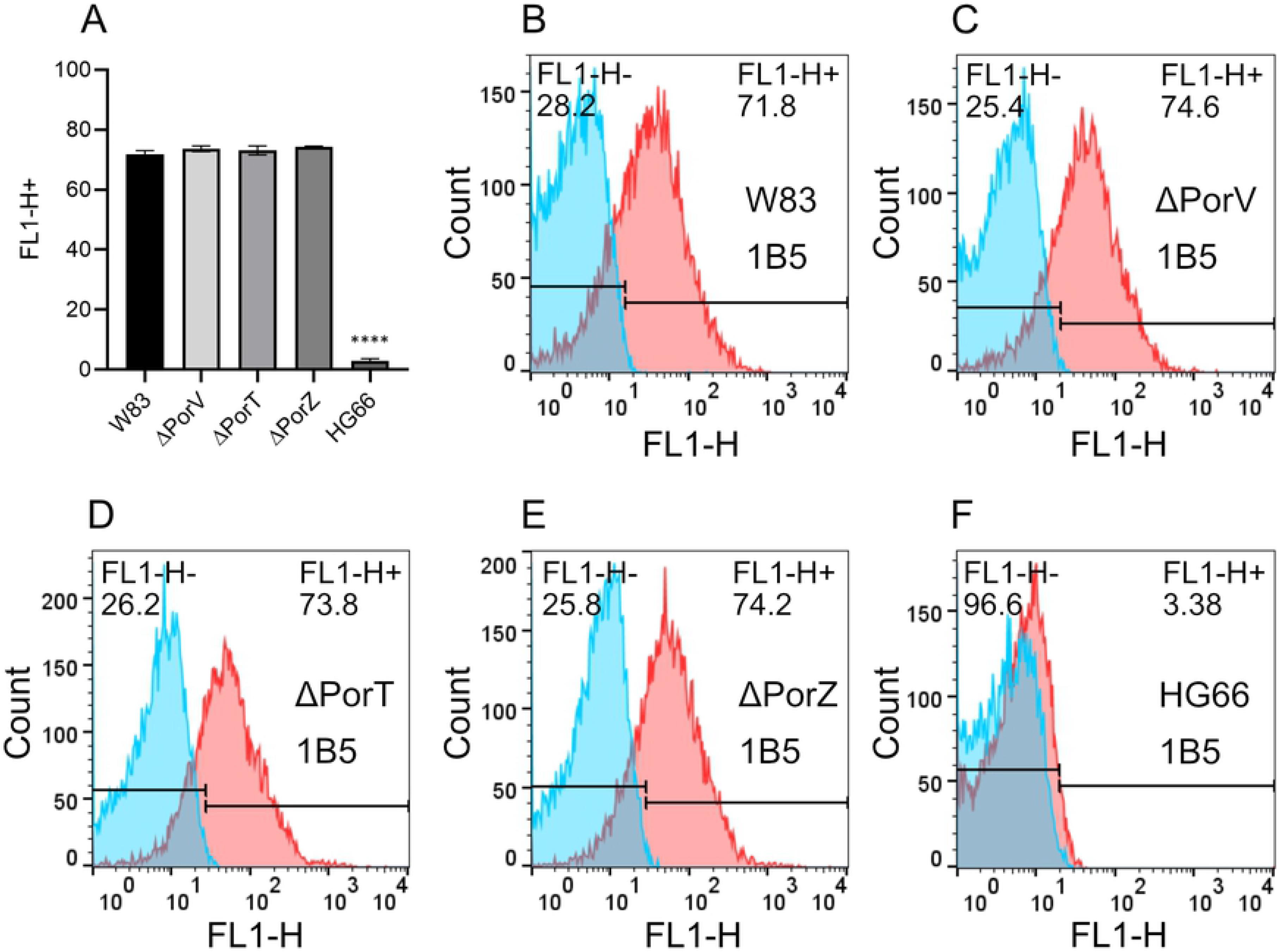
Exposure of A-LPS on the outer membrane of *Porphyromonas gingivalis*. Flow cytometry using mAb 1B5 antibodies to determine surface exposure of A-LPS in ΔPorV, ΔPorT, ΔPorZ mutants, and in W83 and HG66 wild-type strains. (A) A-LPS surface exposure as mean percentage of positive cells in the indicated strains, calculated based on the flow cytometry of three different cultures. (B–F) Representative histograms showing the percentage of mAb 1B5-labelled cells (shown in red) relative to the negative Ab isotype control (shown in blue). Differences between the wild-type W83 parental strain and the secretion mutants were analysed by one-way ANOVA with Bonferroni’s correction; ^****^P < 0.0001.

### PorZ interacts with free periplasmic A-LPS from ΔPorZ, ΔPorV, and ΔPorT mutants

We recently solved the crystal structure of full-length PorZ [32]. The structure revealed that the protein consists of three domains: two seven-stranded β-propeller domains and a C-terminal seven-stranded β-sandwich domain. The latter resembles the canonical CTDs of other T9SS secreted proteins, but in contrast to T9SS cargos, it is not cleaved by PorU. Such β-propeller domains are often engaged in protein–protein and protein–ligand interactions [41]. Furthermore, the architecture of PorZ is similar to that of the periplasmic portion of histidine kinase BT4663 from *Bacteroides thetaiotaomicron*, a most abundant colonizer of the human gut. BT4663 is one part of a two-component system that plays a role in detection and degradation of complex carbohydrates [42]. Based on these structural similarities, we hypothesized that PorZ may also have a glycan-binding function. Specifically, by binding A-LPS, it might be engaged in post-translational modifications of T9SS cargos, resulting in OM anchorage. To test this hypothesis, we investigated whether PorZ could bind the A-LPS in the periplasm of the ΔPorZ, ΔPorV, and ΔPorT mutants. To this end, we treated periplasmic fractions overnight with proteinase K to remove proteins, concentrated them by ultrafiltration, and adjusted them to equivalent LPS content based on the *Limulus* amebocyte lysate (LAL) assay. We then quantitatively analysed PorZ–LPS interaction by microscale thermophoresis. The experiments were performed with fluorescently labelled PorZ and serial dilutions of each periplasmic fraction (Fig 4). As expected, we observed no interaction with components of the HG66 periplasm (Fig 4B) and only weak interaction with W83 (*K*_*d*_ > 10^8^ EU/ml) (Fig 4A). This is consistent with the absence of A-LPS in the cellular compartment of this strain, as determined by western blot analysis (Fig 2B, PP). By contrast, PorZ strongly bound A-LPS from the periplasm of ΔPorZ, ΔPorV, and ΔPorT, with *K*_*d*_ values of 4.01 × 10^5^ EU/ml, 1.51 × 10^5^ EU/ml, and 1.42 × 10^5^ EU/ml, respectively. (Fig 4C, E, G). However, preincubation of the periplasm with mAb 1B5 completely abolished these interactions (Fig 4D, F, H). Given that mAb 1B5 specifically recognizes phosphorylated branched mannan [10], which is a mosaic component of A-PS within A-LPS, these results strongly argue that this unique saccharide moiety is the recognition motif of PorZ.

**Fig 4.**
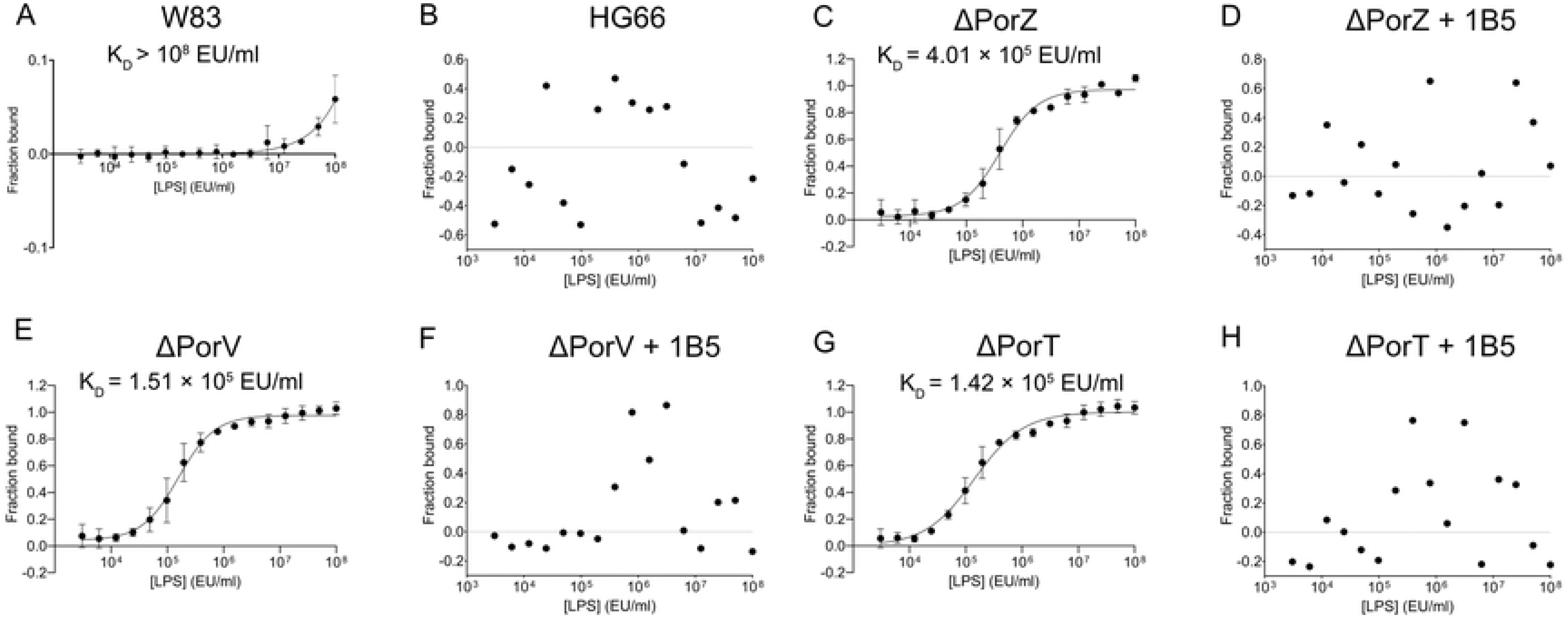
A-LPS in the periplasm of T9SS secretion mutants binds PorZ, and binding is blocked by 1B5 antibodies. Fluorescently labelled PorZ was titrated with serial dilutions of periplasmic fraction from W83 (A), HG66 (B), ΔPorZ (C, D), ΔPorV (E, F), and ΔPorT (G, H) treated overnight with proteinase K and standardized to the same concentration of LPS based on Limulus amebocyte lysate assay, and then subjected to microscale thermophoresis analysis. To confirm the specificity of binding, periplasmic fractions were preincubated with mAb 1B5 (D, F, H). Binding data were curve-fitted and used to determine *K*_*d*_ values. The results are presented as the mean ± SD from three experiments.

### PorZ interacts with purified A-LPS

To verify that PorZ indeed binds the A-LPS, we performed pulldown assays on PorZ in the presence of the A-LPS extracted and purified from whole cells of the ΔPorV mutant and the parental wild-type strain. To this end, we first incubated the A-LPS with immobilised PorZ in an affinity chromatography column, followed by extensive washing and elution of PorZ, and then subjected the eluted fraction to western blot analysis with mAb 1B5. Although the A-LPS from whole ΔPorV and wild-type cells had a wide range of sizes (35–55 kDa and 20–70 kDa, respectively), purified A-LPS migrated more homogeneously (35–40 kDa) (Fig 5A), reminiscent of the A-LPS species from the periplasmic fractions of the secretion mutants (Fig 2B, PP). Notably in this regard, no mAb 1B5-immunoreactive material was eluted from a column that was not charged with PorZ (data not shown).

**Fig 5.**
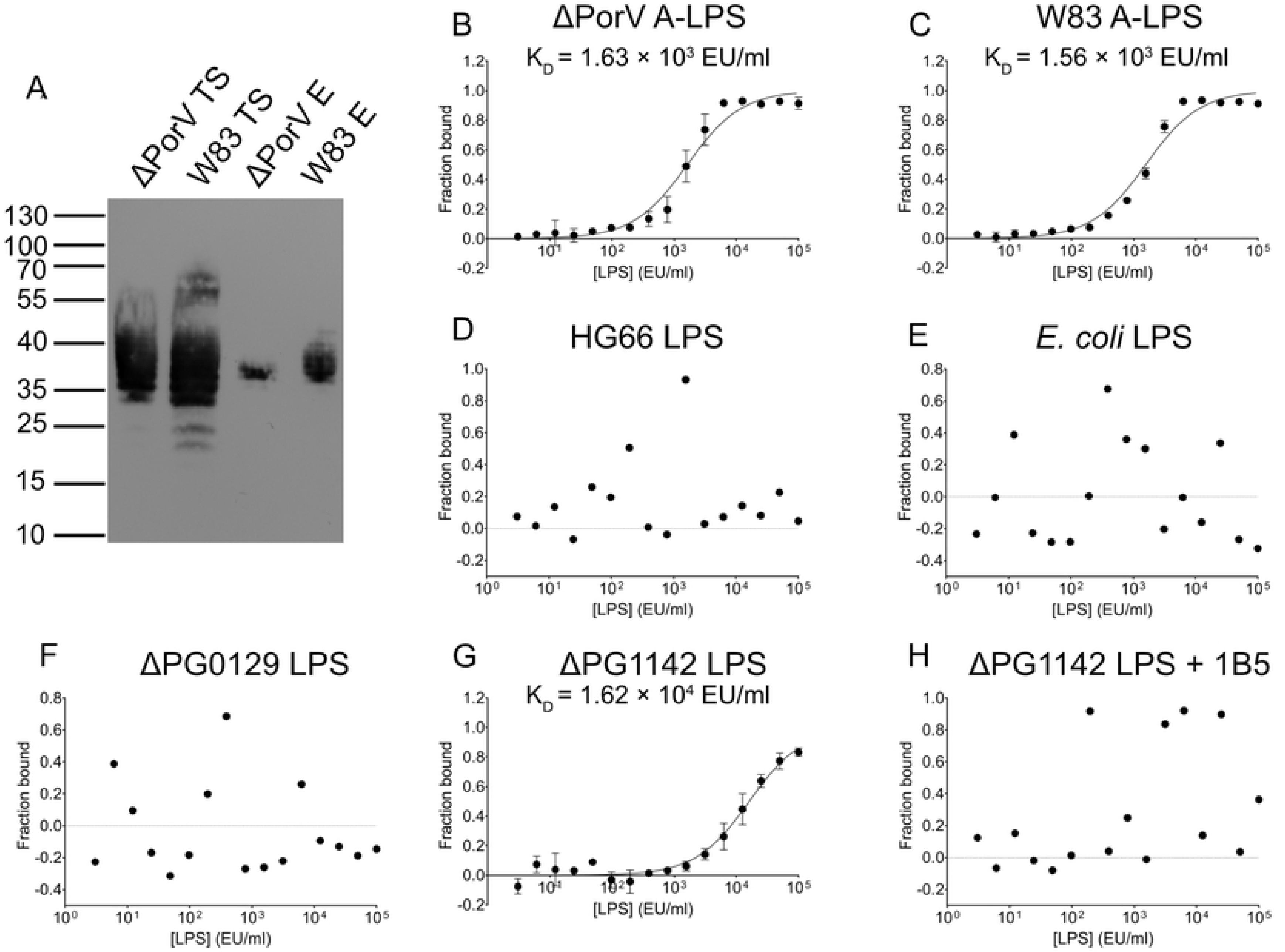
PorZ binds a subset of *Porphyromonas gingivalis* W83-derived A-LPS, dependent on the presence of branched phosphomannan units in A-polysaccharide. (A) GST-tagged PorZ immobilised on glutathione–Sepharose resin was incubated with purified A-LPS from ΔPorV and W83 strains (Loaded A-LPS), and the resin was washed extensively. PorZ, together with interacting A-LPS, was released by cleavage with PreScission protease and analysed for the presence of A-LPS by western blot using mAb 1B5 (PorZ-bound A-LPS). (B) Microscale thermophoresis analysis of fluorescently labelled PorZ incubated with serial dilutions of A-LPS purified from ΔPorV (B) and W83 (C), or total LPS extracted from HG66 (D), *Escherichia coli* (E), ΔPG0129 (F), and ΔPG1142 in the absence (G) and presence of 1B5 (H). Binding data were curve-fitted and used to determine *K*_*d*_ values. The results are presented as means ± SD from three experiments. TS, total sample; E, eluted fraction.

The interaction between PorZ and purified A-LPS was further examined by microscale thermophoresis (Fig 5B–H), as in the aforementioned experiments with periplasm-derived A-LPS. Fluorogenic PorZ was titrated with serial dilutions of A-LPS purified from W83 and ΔPorV. In the case of strain HG66 and mutants ΔPG0129 and ΔPG1142, total LPS (A-LPS and O-LPS) was extracted. The latter two mutants are deficient in α-1,3-mannosyltransferase and O-antigen polymerase activity, respectively [36]. ΔPG0129 synthesizes a deep-R-type LPS with a truncated core region and does not immunoreact with mAb 1B5 [43]. By contrast, ΔPG1142 does immunoreact with mAb 1B5 in western blots because it makes SR-type LPS (core plus 1 repeated unit), containing both an SR-type O-LPS and SR-type A-LPS [43]. The A-LPS extracted from ΔPorV and W83 strains interacted with PorZ with similar affinities (*K*_*d*_ = 1.63 × 10^3^ EU/ml and 1.56 × 10^3^ EU/ml, respectively), apparently because the concentrations of the A-LPS in the two extracts were similar (Fig 5B, C). Notably, the difference in *K*_*d*_ between PorZ-binding periplasm (Fig 4E) and the whole cell-derived A-LPS from the PorV mutant strain (Fig 5B) was the result of normalization of LPS content according to the LAL assay; the periplasm contains a much higher titre of O-LPS than purified A-LPS. By contrast, LPS from HG66 and *Escherichia coli* did not bind to PorZ (Fig 5D, E), further corroborating the finding that regular O-LPS does not interact with PorZ. Remarkably, PorZ-bound A-LPS was produced by the ΔPG1142 strain but not by ΔPG0129 (Fig 5F, G). Because the difference between the A-LPS molecules produced by ΔPG1142 and ΔPG0129 is a single repeat of the phosphorylated branched mannan in SR-type A-LPS produced by the former, but absent in the deep-R-type LPS synthetized by the latter, the phosphorylated branched mannan must be the structure involved in the relatively low-affinity interaction with PorZ (*K*_*d*_ = 1.62 × 10^4^ EU/ml). In keeping with this contention, deep-R-type A-LPS produced by ΔPG0129 did not bind PorZ (Fig 5F).

Moreover, the interaction between ΔPG1142-derived A-LPS and PorZ was completely abolished in the presence of mAb 1B5 (Fig 5H). Together, these results unambiguously argue that PorZ specifically recognizes the phosphorylated branched mannan moiety in A-LPS.

### PorZ preferentially interacts with PorU in the presence of A-LPS

Recently, we showed that surface exposure of PorU is dependent on the presence of PorZ [32]. To determine whether this dependence results from a direct interaction between both molecules, we expressed and purified both proteins and performed co-purification analysis. Most PorZ bound immobilised His-tagged PorU, indicating that the two proteins interact (Fig 6A). To confirm this interaction, we performed microscale thermophoresis analysis. In these experiments, fluorescently labelled PorZ was titrated with serial dilutions of PorU in the absence (Fig 6B) or presence (Fig 6C) of purified A-LPS from W83. In the presence of A-LPS, the affinity of PorZ for PorU increased 5-fold, as indicated by the difference in the respective dissociation constants (*K*_*d*_ = 672 ± 155 nM and 3.38 ± 0.7 μM in the presence and absence of A-LPS, respectively). The stronger binding of A-LPS-loaded PorZ and PorU may play a crucial role in attachment of A-LPS to the C-terminal residue of T9SS cargos during cleavage of the CTD by PorU.

**Fig 6.**
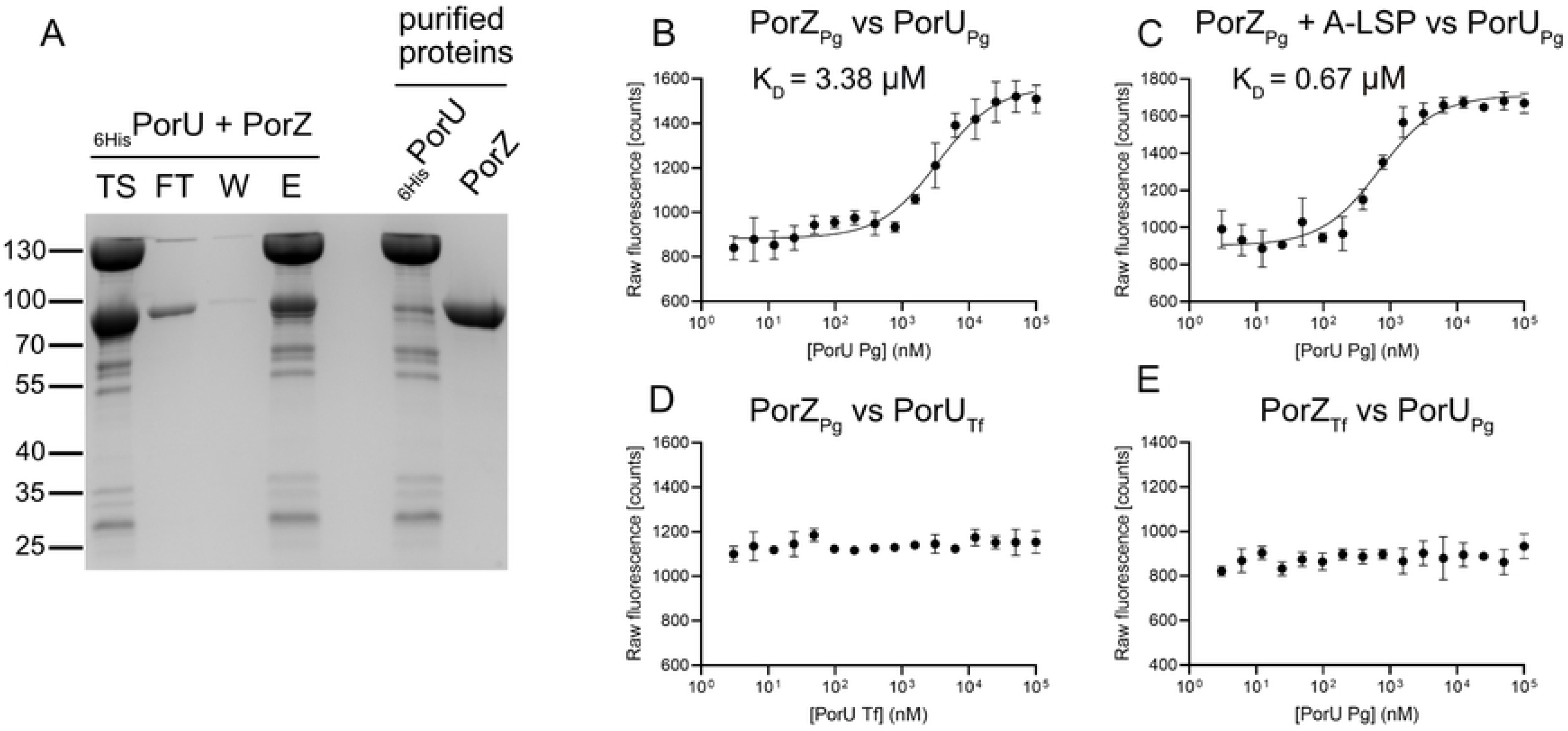
PorZ binds PorU sortase with affinity enhanced by A-LPS. (A) In a pull-out assay, purified recombinant proteins 6His-PorU and PorZ (right two lanes) were mixed and incubated with cobalt-based magnetic beads. The beads were washed, and the bound protein was eluted. The resultant fractions were analysed by SDS-PAGE and Coomassie blue staining. TS, sample before loading on beads; FT, flow-through; W, bead wash; E, eluted proteins. (B–D) Fluorescently labelled PorZ from *Porphyromonas gingivalis* (PorZ_Pg_) was titrated with increasing concentrations of PorU from *P. gingivalis* (PorU_Pg_) without A-LPS (B), in the presence of 1 × 10^5^ EU/ml A-LPS from W83 (C) or titrated with increasing concentrations of PorU from *Tannerella forsythia* (PorU_Tf_). In the analogous experiment, fluorescently labelled PorZ from *T. forsythia* (PorZ_Tf_) was titrated with increasing concentrations of PorU from *P. gingivalis* (PorU_Pg_) (E). Binding data were curve-fitted and used to determine *K*_*d*_ values. The results are presented as means ± SD from three experiments.

### PorU–PorZ interactions are species-specific

T9SS is conserved in many *Bacteroidetes* species, including *Tannerella forsythia*, which is strongly implicated in the pathogenesis of chronic periodontitis [44]. Like *P. gingivalis*, *T. forsythia* secretes many virulence factors via a T9SS and anchors them in the OM [45]. To compare the function of PorU and PorZ in these two periodontal pathogens, we investigated reciprocal interactions between these two proteins in homogenous and heterologous systems. Interestingly, neither *T. forsythia* protein cross-reacted with its *P. gingivalis* ortholog (Fig 6D, E). Although this lack of cross-reactivity can be attributed to the relatively low level of conservation of the primary structures (S4A, B Fig), it is more likely due to specialized functions of the two systems since *T. forsythia* PorZ does not bind A-LPS from *P. gingivalis* (S4C Fig).

### Investigation of the PorZ residues potentially involved in A-LPS binding

The crystal structure of PorZ [32] revealed three positively charged residues exposed to the environment in the β-propeller domains: K42, R132, and R338. These residues are located on the surface of the entry side of the β-propeller domain βD1 (K42 and R132) and the exit side of βD2 (R388) (Fig 7A). Because the A-LPS is negatively charged, we hypothesized that these residues are involved in A-LPS binding by PorZ. To test this hypothesis, we generated the following substitution mutants of PorZ: K42A, R132A, R338A, K42A/R338A, and K42A/R132A/R338A. However, all these mutants exhibited the wild-type phenotype, reflected by black pigmentation (S3A Fig) and production of active gingipains (Fig 7B, C), arguing against the importance of these residues in A-LPS binding.

**Fig 7.**
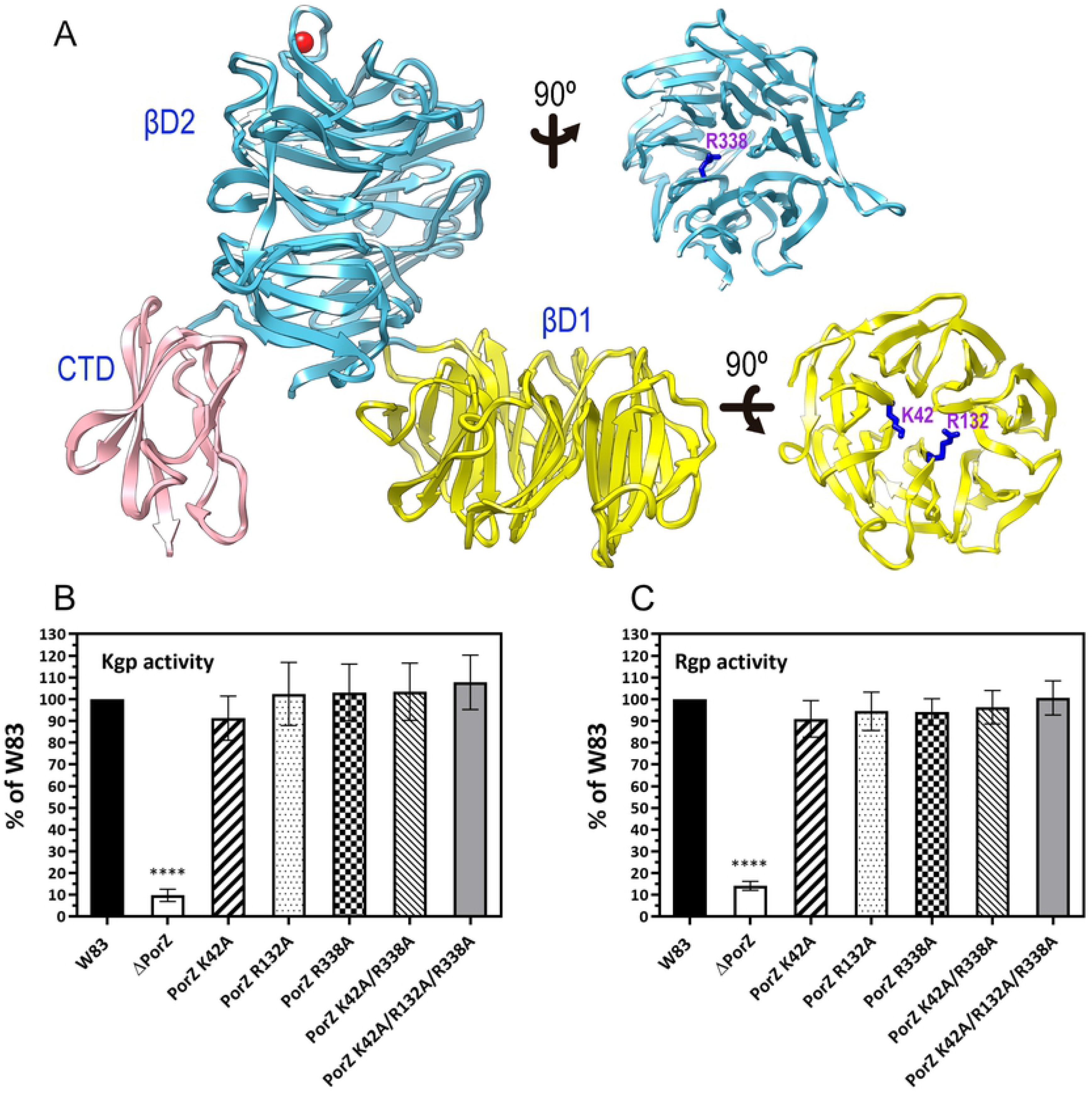
Structure-based analysis of the A-LPS binding site in PorZ. (A) Structure of *Porphyromonas gingivalis* PorZ as a ribbon-type model depicting each of the three constituting domains, β-propeller domains 1 (βD1) and 2 (βD2), and the C-terminal domain (CTD), in one colour. The side chains of three basic residues checked for their role in binding A-LPS (K24, R132, and R338) are shown as dark blue sticks. (B, C) Enzymatic activity of Kgp (B) and RgpA/RgpB (C) in whole cultures as determined with specific synthetic substrates. Cultures were adjusted to OD_600_ = 1.0 prior to testing and processing, and the results shown correspond to triplicate experiments. Significant differences between the wild type and mutants are indicated by *P < 0.05 and ****P < 0.0001.

## Discussion

*P. gingivalis* contains two different lipopolysaccharides (LPS) in its OM, O-LPS and A-LPS [9]. The main difference between them is limited to the chemical composition of the O-antigen. The tetrasaccharide repeat unit in O-LPS consists of a conventional polysaccharide (PS), which is distinct from the anionic polysaccharide (APS) repeat unit in A-LPS (Fig 1). The APS unit contains a phosphorylated branched mannan recognized specifically by mAb 1B5 [10]. This serves as a convenient tool for visualizing A-LPS alone or conjugated to T9SS cargo proteins by a variety of immunological methods [33]. Using this approach, we showed that *P. gingivalis* mutants with dysfunctional T9SS accumulated A-LPS in a protein-unconjugated form in the periplasm, regardless of which component of the T9SS machinery was inactivated by mutagenesis (Fig 2B, PP).

A-LPS accumulated in the periplasm of ΔPorZ, ΔPorV, and ΔPorT strains, and interacted with PorZ. The dissociation constant (*K*_*d*_), determined by microscale thermophoresis (MST), differed from sample to sample, suggesting variations in A-LPS concentration (ΔPorT > ΔPorV > ΔPorZ) in the periplasm of these secretion mutants (Fig 4). The *K*_*d*_ calculation was based on the concentration of total LPS estimated using the LAL assay, with samples adjusted to the same titre of LPS regardless of the A-LPS:O-LPS ratio. Consequently, the values obtained are only semi-quantitative. Nevertheless, the general usefulness of the assay was validated by the fact that PorZ did not interact with the periplasm derived from strain HG66, which produces intact O-LPS but lacks A-LPS [39], and interacted very weakly with periplasm derived from wild-type *P. gingivalis*. These observations correlate perfectly with the absence of immunoreactive A-LPS in these fractions (Fig 2). In the latter case, the interactions detected by MST were probably due to periplasmic contamination with OM-derived A-LPS acquired during subcellular fractionation.

A previous study reported that PorV has an LPS deacylase activity, and the ΔPorV mutant produces only penta-acylated forms of mono-phosphorylated lipid; by contrast, wild-type *P. gingivalis* and the ΔPorT mutant both have tetra- and penta-acylated forms of mono-phosphorylated lipid A [33]. The comparable *K*_*d*_ values of PorZ interactions with A-LPS in periplasm derived from these strains clearly suggests that even if PorV has deacylase activity, which was questioned by a subsequent study [46], the acylation profile of lipid A has no influence on this binding. Also, differences in lipid A phosphorylation, which appear to be influenced by PorV [46], did not have any effect on the interaction between A-LPS and PorZ, as the *K*_*d*_ values were comparable for all tested secretion mutants. Hence, A-LPS binding to PorV may be independent of the structure of lipid A.

Together, these observations indicate that A-LPS, regardless of its lipid A acylation and/or phosphorylation, binds specifically to PorZ. The presence of A-LPS in the periplasm of secretion mutants was in stark contrast to the wild-type strain, which almost completely lacked A-LPS in the periplasm. Remarkably, periplasmic accumulation of A-LPS in secretion mutants did not affect its level in the OM (Fig 3). This is intriguing in the context of LPS biosynthesis and transport from the cytoplasm across the cellular envelope to the bacterial surface.

The lipid A-core oligosaccharide complex is synthesized on the cytoplasmic face of the IM and flipped to the periplasmic side of the IM [47]. Here, O-antigen, which is independently synthesized in the cytoplasm and translocated to the periplasmic face of the IM as undecaprenol phosphate-linked O-antigen (in *P. gingivalis*, A-PS, or O-PS), is ligated to lipid A-core oligosaccharide by the IM-associated WaaL ligase. Then, the mature LPS (in *P. gingivalis* O-LPS and A-LPS) is transported to the cell surface by the Lpt (<u>LPS</u> transport) pathway, which involves several essential proteins that bridge IM and OM in the periplasm [48]. Notably in this regard, all Lpt components are conserved in *P. gingivalis*. The transport occurs in a continuous manner (the PEZ model), without any LPS leaking into the periplasm [40]. Therefore, the presence of considerable amounts of A-LPS in the periplasm of *P. gingivalis* T9SS mutants is surprising because it suggests the existence of a selective shunt of A-LPS into the periplasm from the continuous flow of the molecules along the Lpt periplasmic bridge. The distinct size of A-LPS in the periplasm (Fig 2A, PP) and its interaction with PorZ (Fig 5A), in contrast to A-LPS extracted from the whole cells (Fig 2A, WC), argues that two distinct pools of A-LPS exist. It is tempting to speculate that this is related to additional modification of A-LPS that is ultimately attachment to secreted proteins via the 648-Da linker [49]. The linker may provide an amino group for nucleophilic attack on the thioester intermediate in the PorU-catalysed proteolysis of a peptide bond that leads to removal of the CTD and attachment of A-LPS to the C-terminus of a truncated T9SS cargo protein [49]. Because PorU is localized on the bacterial surface, this transpeptidation reaction must occur after A-LPS, with its linker, translocates to the OM, where it encounters T9SS cargo proteins and PorU. In T9SS mutants, translocation is apparently dysfunctional, leading to accumulation of A-LPS in the periplasm.

An elegant recent study by the Reynolds showed that the surface-localized (PorU and PorZ) and integral OM β-barrel proteins (PorV and PorQ) form a 440 kDa attachment complex [29]. In their model, PorV serves to shuttle T9SS cargo proteins from the OM translocon to the attachment complex. There, PorU sortase cleaves the CTD and attaches A-LPS provided by PorZ to T9SS cargo proteins, thereby facilitating anchorage to the OM. Our results experimentally validate this model by showing that A-LPS does indeed interact with PorZ. The signature pattern recognized by PorZ was unambiguously mapped to a phosphorylated branched mannan (Manα1-2Manα1-phosphate) in APS (A-PS) (Figs 4 and 5). A single repeat unit of A-LPS is sufficient to interact with PorZ, but with decreased affinity (Fig 5G) that is not sufficient to mediate attachment of A-LPS to secreted proteins. The *P. gingivalis* ΔPG1142 mutant strain, which produces severely truncated A-PS carrying a single repeat of the phosphorylated branched mannan, is unable to retain gingipains on the cell surface and instead releases them in unmodified, soluble forms into the medium [36].

The A-PS binding site of PorZ has yet to be determined. The mutation of Arg residues in the β-propeller domains of PorZ did not affect the *P. gingivalis* secretory phenotype (S3A Fig). Interestingly, however, we showed for the first time that PorZ directly interacts with PorU, and that this interaction was significantly strengthened in the presence of A-LPS (Fig 6B, C). This change in the affinity may be essential for A-LPS transfer from PorZ to T9SS client proteins mediated by the sortase activity of PorU. Once CTD is cleaved and released into the growth medium [30] and A-LPS is attached to a cargo protein, both PorV and PorZ are recycled for another round of coordinated transport and attachment.

Taken together, our findings not only complement the previously proposed model of molecular events occurring on the *P. gingivalis* surface during protein secretion through T9SS [29], but also add a significant mechanistic insight. First, our data argue that A-LPS designated for attachment to T9SS cargo proteins is probably transported to the cell surface via a periplasmic/OM shunt independent of the Lpt pathway of O- and A-LPS insertion into the outer layer of the OM. Second, PorZ functions as the A-LPS shuttle protein, specifically recognizing the phosphorylated branched mannan in the repeat units of the A-PS. Third, enhanced affinity of A-LPS loaded PorZ for the PorU facilitates presentation of A-LPS to the sortase. The sortase cleaves the CTD from the T9SS client protein and simultaneously catalyses a transpeptidation reaction that attaches A-LPS to the C-terminal carbonyl group of the secreted protein via an isopeptide bond [49].

## Supporting information

**S1 Fig. A-LPS is conjugated to cargo proteins only in the wild-type strain W83.**

(A, B) Proteinase K treatment of periplasmic fraction (A) and washed cells (B) of *Porphyromonas gingivalis* mutants and wild-type strain W83. The fractions were incubated at 60°C for 1 h, treated overnight with proteinase K, and then subjected to western blot with mAb 1B5. PK, proteinase K.

**S2 Fig. Schematic view of the strategy for *Porphyromonas gingivalis* point mutant generation.**

**S3 Fig. Phenotypes of PorZ mutants.**

Mutants were grown on eTSB blood agar plates for 3 days.

1. ΔPorZ; 2. W83; 3. PorZ K42A; 4. PorZ R132A; 5. PorZ R338A; 6. PorZ K42A/R338A; 7. PorZ K42A/R132A/R338A.

**S4 Fig. Sequence alignments for PorZ (A) and PorU (B) from *Porphyromonas gingivalis* W83 and *Tannerella forsythia.*** The secondary structure assignment, based on the crystal structure of PorZ from *P. gingivalis*, is indicated. The image was produced by ESPript 3.01 [50]. Sequence alignment was performed using Clustal Omega [51]. (C) Specificity of PorZ from *P. gingivalis*. Fluorescently labelled PorZ from *T. forsythia* was titrated with increasing concentrations of purified A-LPS from W83. The results are presented as means ± SD from three experiments.

